# Association of poultry vaccination with the interspecies transmission and molecular evolution of H5 subtype avian influenza virus

**DOI:** 10.1101/2023.12.20.572711

**Authors:** Bingying Li, Jayna Raghwani, Sarah C. Hill, Sarah François, Noémie Lefrancq, Yilin Liang, Zengmiao Wang, Lu Dong, Phillipe Lemey, Oliver G. Pybus, Huaiyu Tian

## Abstract

The effectiveness of vaccinating poultry in preventing the transmission of highly pathogenic avian influenza viruses (AIVs) has been questioned for years and its impact on wild birds is uncertain ^1–3^. Here we reconstruct movements of H5 subtype AIV lineages among vaccinated poultry, unvaccinated poultry, and wild birds, worldwide from 1996 to 2023. We find that lineage transitions among host types are lagged and that movements from wild birds to unvaccinated poultry were more frequent than those from wild birds to vaccinated poultry. However, we also find that the HA gene of the AIV lineage that circulated predominately among Chinese poultry with high vaccination coverage underwent faster evolution and greater nonsynonymous divergence than other lineages. Further, this Chinese poultry lineage contained more codons inferred to be under positive selection, including at known antigenic sites, and its rates of nonsynonymous divergence and adaptative fixation increased after mass poultry vaccination began. Our results indicate that the epidemiological, ecological and evolutionary consequences of widespread AIV vaccination in poultry may be linked in complex ways, and that much work is needed to better understand how such interventions may affect AIV transmission to, within and from wild birds.

## Introduction

During the summer of 2022, seabirds in many European, North American, and African countries suffered unprecedented mortality from avian influenza virus (AIV) ^4^. The causative virus, a highly pathogenic H5 subtype AIV (HPAIV) belonging to clade 2.3.4.4b, has been detected at unprecedented incidence in wild birds ^5^. Wild birds, the natural reservoir of AIVs, can acquire and transmit viruses to poultry or mammals ^6, 7^ and play a major role in the maintenance and global dissemination of AIVs ^8^. Intra- and inter-species transmission of genetically diverse AIVs can result in virus genomic reassortment ^9, 10^ and the emergence of novel HPAIV lineages ^11–19^.

In order to protect poultry from infection with HPAIV, some countries have implemented vaccination programs in poultry for H5 subtype avian influenza, mostly in Asia and Africa (Fig. 1) ^20–23^. Based on national vaccination data from 2010, Egypt had the highest vaccination coverage in poultry (82%), followed by China (73% in 2009 and 87% in chickens and < 30% in ducks in 2018 ^24^), Vietnam (31%), and Indonesia (12%) ^25^. In 2013, vaccination coverage for commercial layer flocks in two districts of Bangladesh were reported to be 32% and 54% ^26^. France and the Netherlands have also vaccinated poultry; however, their overall vaccination coverages are very low (<0.1%). Since 2005 China has implemented a nationwide vaccination program ^27^ and notably accounts for >90% of the global consumption of H5 AIV vaccines. Due to the continuing evolution of H5 AIV, vaccine strains are frequently updated to ensure their effectiveness ^28, 29^. Several studies have suggested that the extensive vaccination of Chinese poultry against H5 AIV has suppressed outbreaks effectively and substantially decreased the prevalence of H5 AIV in live bird markets ^26, 29–32^.

**Fig 1.**
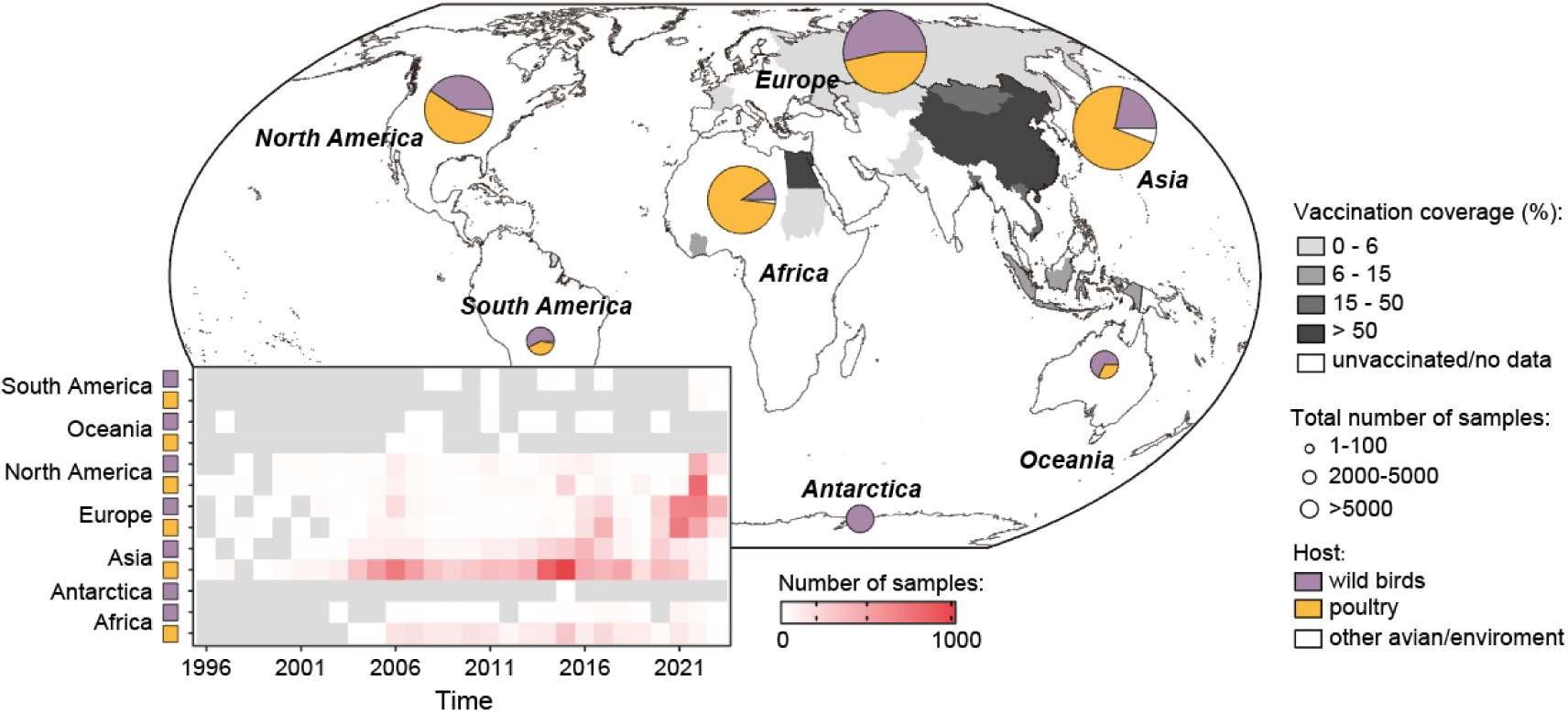
**Vaccination coverage and number of H5 AIV HA gene sequences across different continents and countries.** Countries are shaded according to the vaccination coverage in poultry in 2010. The pie charts show the total number of H5 AIV HA gene sequences sampled from poultry, wild birds, and environment since 1996. Viral sequences are categorized according to their host: wild birds (purple), poultry (yellow), and other avian/environment (white). Inset: number of H5 AIV HA gene sequences sampled in poultry and wild birds, per year, per continent.

However, there are concern that mass vaccination against H5 AIV could impact the molecular evolution of the virus ^33^. For example, after mass vaccination was initiated against H5N1 AIV in China, a significant increase in the evolutionary rate of H5N1 AIV was observed during 2005-2010 ^34^. Similarly, in North America, the evolutionary rate of H5N2 AIV in Mexico was inferred to be significantly higher between 1993 and 2002, following a period of mass avian influenza vaccination in this region, compared to the rate of evolution of H5 AIV in the United States, where vaccination was not used ^35^.

Furthermore, AIV lineages circulating in Egypt and Indonesia, where vaccination against H5N1 is prevalent, are characterized by higher evolutionary rates and a greater number of positively selected sites in the HA gene compared to countries where vaccination is about, such as Nigeria, Turkey and Thailand ^36, 37^. However, these studies have concentrated mainly on viruses from poultry and lacked data from wild birds and at the wild bird-poultry interface. This limitation could potentially create confusion in understanding the evolutionary characteristics of the virus in both wild birds and poultry, contributing to the uncertainty of their conclusions. Given the frequent movement of AIV between wild birds and poultry, it is crucial to understand the impact of mass poultry vaccination on AIV in unvaccinated wild birds.

Here we conduct phylogenetic analyses to investigate the inter-species transmission of H5 AIV between wild birds and poultry from 1996 to 2023, and compare the evolutionary dynamics of H5 AIV in different host populations that vary in vaccination status.

## Result

### Interspecies transmission of H5 AIV at the interface of poultry and wild birds

We collated a total of 22,606 hemagglutinin (HA) gene sequences belonging to H5 AIV, sampled from each continent since 1996 (Fig. 1). The viral sequences were unevenly distributed over time and space across host populations. Therefore, for the downstream analyses, we focused on Europe and Asia, where sufficient viral genetic data was available from long-term sampling of poultry and wild birds to quantify the dynamics of virus transmission and evolution within and among host populations. By taking into account the implementation of vaccination programs and the availability of viral genetic sequences (Supplementary Fig. S1), we categorised sequences into eight groups based on the country of sampling, host species, and vaccination status: wild birds (without vaccination), European poultry (without vaccination), Japanese poultry (without vaccination), Korean poultry (without vaccination), Indonesian poultry (with vaccination, low vaccination coverage), Bangladeshi poultry (with vaccination, low vaccination coverage), Vietnamese poultry (with vaccination, low vaccination coverage), and Chinese poultry (with vaccination, high vaccination coverage).

To investigate H5 AIV transmission and evolution, we estimated a time-resolved phylogeny and inferred ancestral state (Fig. 2A; Supplementary Table S1). Inter- species movement of H5 AIV lineages from Chinese poultry to wild birds, and from wild birds to European poultry, were frequently observed (Fig. 2B). Bayesian reconstruction of the host status of H5 AIV lineages indicates there were three waves of spillovers from Chinese poultry to wild birds (CPtoWB), and from wild birds to European poultry (WBtoEP) (Fig. 2C). To investigate lineage transitions among these three host types before 2020, we conducted an extended convergent cross mapping (CCM) analysis on the time series of mean annual lineage transitions (Markov jumps) from Chinese poultry to wild birds, and from wild birds to European poultry, between 1996 and 2019. We find that CPtoWB transitions preceded WBtoEP transitions, with a time lag of approximately 2-3 years (Fig. 2C; Supplementary Fig. S2). The observed pattern is robust to diverse genomic sampling strategies (Supplementary Figs. S3-S6).

**Fig 2.**
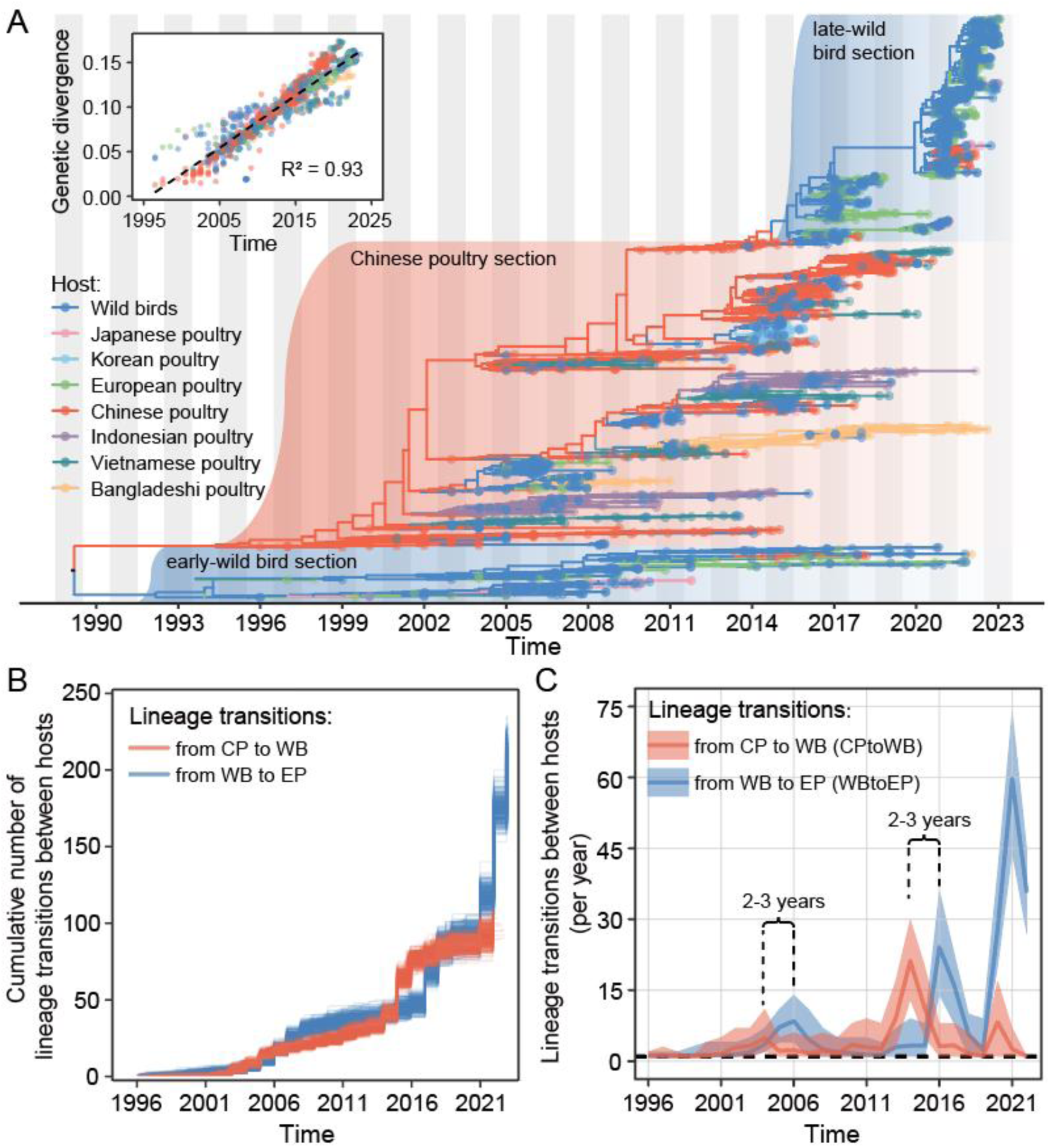
**Temporal dynamics of H5 AIV lineage transitions between wild birds and poultry.** (**A**) The maximum clade credibility tree of H5 AIV sequences sampled from 1996 to 2023. Tree tips are coloured according to the host from which the sequence was sampled, while internal branches represent ancestral host states inferred using the asymmetric discrete phylogenetic model (dark blue: wild birds; orange: Chinese poultry; light green: European poultry; purple: Indonesian poultry; pink: Japanese poultry; light blue: Korean poultry; dark green: Vietnamese poultry; yellow: Bangladeshi poultry). Inset: a root-to-tip regression of genetic divergence against dates of sample collection. (**B**) Cumulative number of host population changes (Markov jumps) on lineages in the HA gene phylogeny. The lineage transitions between hosts were summarized from a posterior sample of trees from the asymmetric discrete phylogenetic model. (**C**) Time series of the annual mean number of HA gene lineage transitions between wild birds, Chinese poultry and European poultry.

Overall, more viral lineage transitions were observed from wild birds to unvaccinated poultry populations than to vaccinated poultry populations (Fig. 3, Supplementary Table S2). Frequent lineage transitions were observed from wild birds to unvaccinated European, Japanese and Korean poultry, especially after 2020 (Fig. 3A). Conversely, there were fewer viral lineage transitions from wild birds to Chinese poultry and other vaccinated poultry populations (Fig. 3B). Inter-species transitions from poultry to wild birds were frequent after 2016 (Fig. 3C and Fig. 3D). Virus lineage movements between poultry populations were not common. The observed pattern is robust to diverse genomic sampling strategies (Supplementary Table S3).

**Fig 3.**
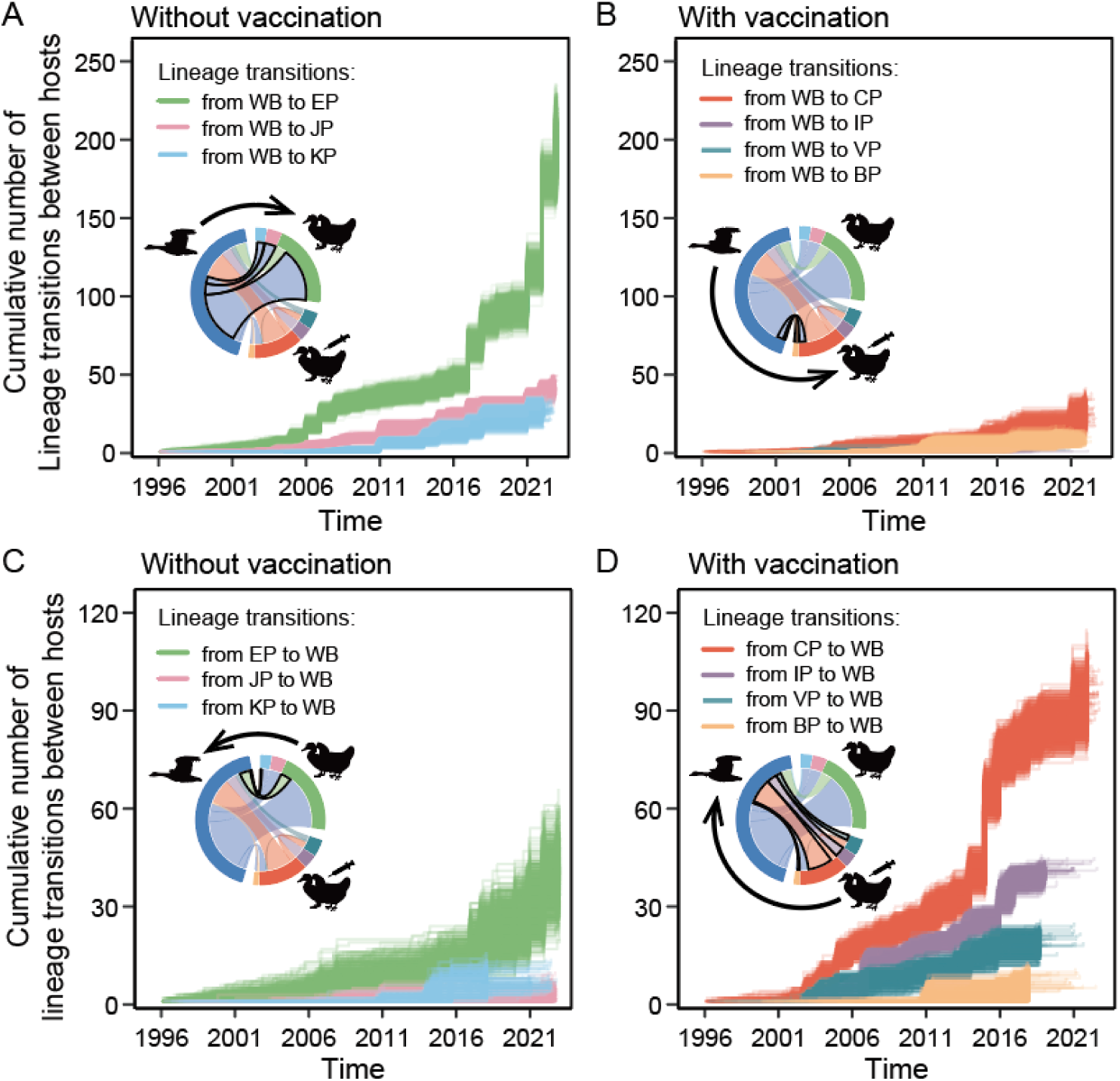
**Inter-species lineage transmission between wild birds and poultry populations with different vaccination statues.** (**A**) Accumulation of lineage transitions from wild birds to unvaccinated poultry populations. (**B**) Accumulation of lineage transitions from wild birds to vaccinated poultry populations. (**C**) Accumulation of lineage transitions from unvaccinated poultry populations to wild birds. (**D**) Accumulation of lineage transitions from vaccinated poultry populations to wild birds. Chord diagrams show the mean cumulative lineage transitions between different groups of sequences (dark blue: wild birds; orange: Chinese poultry; light green: European poultry, purple: Indonesian poultry; pink: Japanese poultry; light blue: Korean poultry; dark green: Vietnamese poultry; yellow: Bangladeshi poultry). Animal silhouettes are from PhyloPic.org. The plots were summarized from a posterior sample of trees from the asymmetric discrete phylogenetic model.

### Evolutionary dynamics of H5 AIV in different host populations

We next investigated the evolutionary dynamics of H5 AIV in wild birds and poultry populations with different vaccination levels. Poultry-dominated viral lineages were identified only for China (vaccinated since 2005), Bangladesh (vaccinated since 2012) and Indonesia (vaccinated since 2004) (Fig. 4A). Our results indicated that the Chinese poultry lineage had a significantly higher substitution rate (mean rate = 5.38×10^-3^ sub/site/year; 95% HPD: 5.02-5.76×10^-3^; *P* < 0.05) than the early-wild bird lineage (3.39×10^-3^ sub/site/year; 95% HPD: 2.97-3.83×10^-3^) (Fig. 4B). Notably this result is robust to the time-dependence of virus evolutionary rates ^38^ because both lineages have similarly high variation in sequence sampling dates (Fig. 4C). Besides, the substitution rate of the Chinese poultry lineage during the vaccination period (5.12×10^-3^ sub/site/year; 95% HPD: 4.71-5.55×10^-3^) was faster than before vaccination (4.79×10^-3^ sub/site/year; 95% HPD: 4.38-5.18×10^-3^; *P* < 0.05; Supplementary Fig. S7). The substitution rates of poultry-dominated viral lineages in Bangladesh and Indonesia were also faster than those of the early-wild bird lineage (*P* < 0.05). Furthermore, the substitution rate of the late-wild bird lineage was faster than that of the Chinese poultry lineage from which it emerged (5.91×10^-3^ sub/site/year; 95% HPD: 5.03-6.56×10^-3^), however this latter result should be interpreted with caution due to the comparatively shorter range of sequence sampling dates in the late- wild bird lineage.

**Fig 4.**
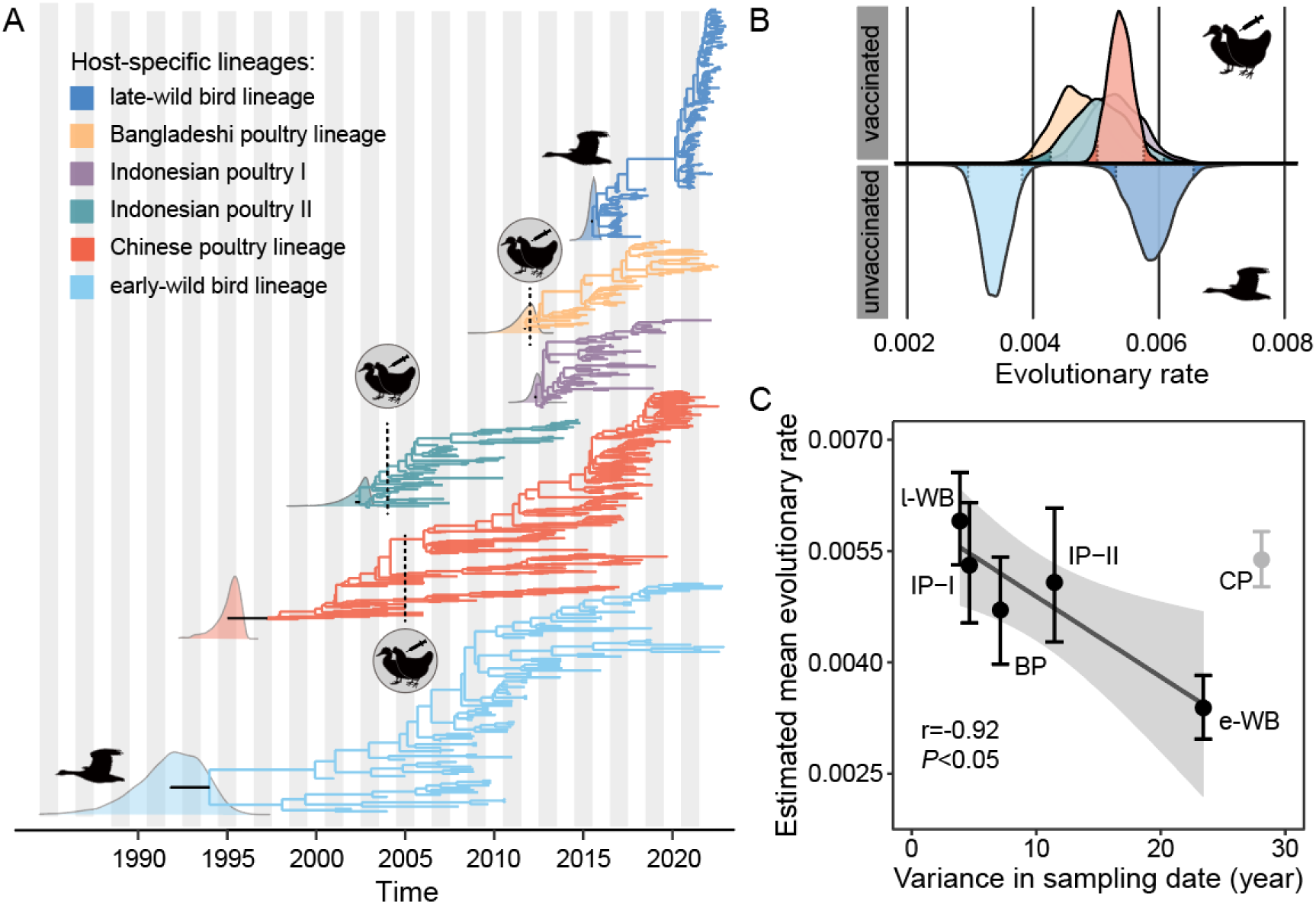
**Evolution of the hemagglutinin (HA) gene of H5 AIV in different host- specific lineages.** (**A**) Time-resolved phylogenies of H5 AIV lineages in wild birds and vaccinated poultry. The dashed line represents the date when each country started implementing avian influenza vaccination for poultry. The density plot shows the estimated tMRCA for each lineage. (**B**) The estimated substitution rates of the HA gene vary among host-specific H5 AIV lineages. Lineages in unvaccinated host populations (the early- and late-wild bird lineages) are shown below the line and those in vaccinated host populations (the Chinese, Bangladeshi and Indonesian poultry lineages) are shown above the line. Highlighted region shows 95% confidence intervals. (**C**) Scatterplot of the variance in sampling date versus estimated evolutionary rate, for each host-specific lineage. The error bars show the 95% confidence intervals for estimated evolutionary rates. A regression analysis (excluding the Chinese poultry lineage data point) was conducted to show the negative relationship between expected due to the time-dependency of inferred evolutionary rates (r = -0.92; *P* < 0.05), and highlight that the Chinese poultry data point is an outlier.

If the higher evolutionary rate of HA gene in vaccinated poultry lineages is simply a result of increased viral transmission facilitated by high poultry densities, then we should also observe higher rates for poultry lineages in other AIV segments.

Conversely, PB2 gene evolution was faster in the wild bird lineage (3.62×10^-3^ sub/site/year; 95% HPD: 3.23-3.92×10^-3^; *P* < 0.05) than in the Chinese poultry lineage (3.44×10^-3^ sub/site/year; 95% HPD: 3.08-3.78×10^-3^; Supplementary Fig. S8- S9). This result suggests that the faster evolution of the HA gene in Chinese poultry may not be attributable to higher transmission rates alone and instead could result from selection on the HA gene.

### Viral adaptive evolution in host populations with different vaccination status

We then investigated whether the higher rate of molecular evolution observed in the Chinese poultry lineage, compared to the early-wild bird lineage, could be explained by greater viral adaptive evolution in poultry. For each lineage, we calculated the nonsynonymous and synonymous divergence of the HA gene through time, from a known reference sequence (NCBI Reference Sequence: AF144305; Figs. 5A, 5B and 5C). Our results indicate higher nonsynonymous divergence for the Chinese poultry lineage than for other lineages, such as wild bird lineages and poultry lineages in Bangladesh and Indonesia (Chinese poultry lineage gradient = 0.0027, *P* < 0.05; early-wild bird lineage = 0.0005, *P* < 0.05; late-wild bird lineage = 0.0007, *P* < 0.05; Bangladeshi poultry lineage = 0.0009, *P* < 0.05; Indonesian poultry lineage I = 0.0011, *P* < 0.05; Indonesian poultry lineage II = 0.0021, *P* < 0.05). Additionally, the nonsynonymous divergence rate increased in the Chinese poultry lineage following mass vaccination (gradient pre-2005 = 0.0017, 95% CI = 0.0011-0.0023; gradient between 2005 and 2010 = 0.0046, 95% CI = 0.0041-0.0052; Supplementary Table S4). To test for the potential impact of different sampling intensities among countries, the Chinese poultry lineage was resampled and the results remained consistent (Supplementary Fig. S10). This indicates that the Chinese poultry lineage exhibited more nonsynonymous evolution, especially after vaccination, compared to the wild bird lineages and the poultry lineages that experienced lower vaccination coverage.

**Fig 5.**
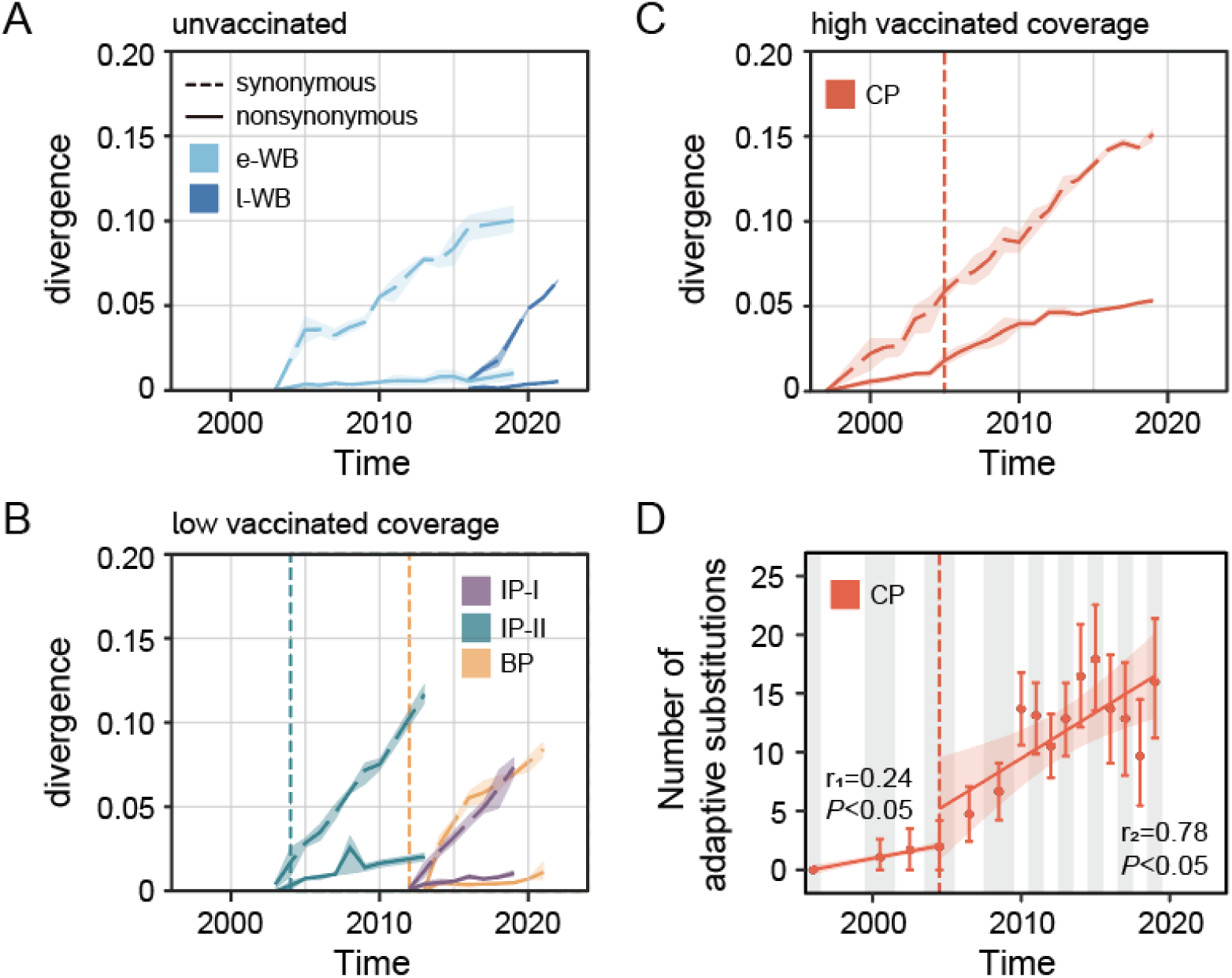
**Temporal dynamics in divergence and adaptive fixation of the H5 AIV HA gene, in different host-specific lineages.** (**A-C**) Nonsynonymous (solid lines) and synonymous (dashed lines) divergence of the HA gene through time. The host-specific lineages were classified into three groups according to the vaccination state: unvaccinated group: (**A**) early-wild bird lineage (e-WB, only the top lineage was retained), late-wild bird linage (l-WB); (**B**) low vaccinated coverage group: Indonesian poultry lineage I (IP-I), Indonesian poultry lineage II (IP-II), Bangladeshi poultry lineage (BP); and (**C**) high vaccinated coverage group: Chinese poultry lineage (CP). Divergences were computed using 1-year sliding windows. Shaded regions show 95% confidence intervals. The dashed line represents the date when each country started implementing avian influenza vaccination for poultry. (**D**) The accumulation of viral adaptative substitutions in the Chinese poultry lineage. Two regression lines were estimated, before (r_1_) and after (r_2_) 2005. The first sequence, sampled in 1996, was used as the ancestral sequence, those sampled from 1997 to 1999 were excluded due to insufficient sample size.

Since nonsynonymous divergence can result from either positive selection or random genetic drift, we also used an independent population genetic method to estimate the rate of accumulation of adaptive substitutions in the HA gene of the Chinese poultry lineage. The estimated rate of adaptative fixation in that lineage was significantly higher after the initiation of vaccination in poultry (gradient pre-2005 = 0.24 adaptive fixations per codon per year; *P*<0.05; gradient after 2005 = 0.78 adaptive fixations per codon per year; *P* < 0.05; Fig. 5D). These estimated adaptation fixation rates are rapid, but lower than those previously inferred for the HA gene of human influenza subtype H3N2 (1.52) and subtype H1N1 (1.02) ^39^. We next estimated dN/dS values for each host-associated lineage using the renaissance counting method ^40^. Mean dN/dS is higher in the Chinese poultry lineage (0.24) than in the early wild bird lineage (0.16), the late-wild bird lineage (0.18), and the European poultry lineage (0.16) (Supplementary Table S5). The mean dN/dS values of the Bangladeshi (0.17) and Indonesian poultry I and II lineages (0.18 for both) were also lower than that of the Chinese poultry lineage.

Furthermore, dN/dS values were calculated for individual codons in the HA gene, for each host-specific H5 AIV lineage. We employed multiple methods to detect sites under positive selection (Table 1). Our results indicate the number of amino acid sites exhibiting evidence of positive selection in the Chinese poultry lineage was greater than in the two wild bird lineages and in the Bangladeshi and Indonesian poultry lineages (Table 1). More positively selected sites were also observed in resampled- Chinese poultry lineage (Supplementary Table S6), indicating the finding is robust to diverse sampling strategies (Supplementary Tables S7, S8).

**Table 1.** Positively-selected sites in the HA gene among different host-specific H5 AIV lineages.

| Host-specific lineages | Vaccination coverage | Livestock density in 2015 (birds/km<sup>2</sup>) † |  | Methods |  |  |  |
| --- | --- | --- | --- | --- | --- | --- | --- |
|  |  | chicken | duck | RC | FEL (*P* < 0.1) | SLAC (*P* < 0.1) | FUBAR (PP > 0.9) |
| Chinese poultry lineage | 73%<sup>25</sup> | 921 | 126 | 3, 61, 142, 156, 157, 171, 172, 178, 185, 205, 242, 285, 289, 527 | 87, 142*, 154*, 156, 157*, 172*, 185, 285*, 291*, 325* | 3, 142*, 154*, 157*, 172*, 185*, 285* | 142*, 154*, 157*, 171, 172*, 285* |
| Bangladesh poultry lineage | <50%<sup>26</sup> | 1448 | 292 | 170, 204, 205 | 204*, 205* | 204, 205 | 170, 204*, 205* |
| Indonesia poultry lineage I | 12%<sup>25</sup> | 1262 | 106 | 170, 204, 211 | 205*, 554 | 205 | 170*, 205* |
| Indonesia poultry lineage II |  |  |  | 10, 156, 529 | 3, 11, 156*, 529 | 156* | 5*, 11*, 156* |
| early-wild bird lineage | - | - |  | 10, 102, 170, 171 | 102*, 170*, 171 | 102*, 170 | 102*, 170*, 171 |
| late-wild bird lineage |  |  |  | 10 | 185, 507 | - | 252 |

†Mean livestock density was calculated using data from the Gridded Livestock of the World (GLW)<sup>80</sup> website, hosted by Food and Agriculture Organization (FAO; https://dataverse.harvard.edu/dataverse/glw). Regions with <10 birds per square kilometer were excluded from the calculation. \*Asterisks mark sites inferred to be under positively selected with posterior probability (PP) >0.95 and <0.05. FEL, Fixed Effects Likelihood. SLAC, Single Likelihood Ancestor Counting. FUBAR, Fast Unconstrained Bayesian AppRoximation. RC, renaissance counting method implemented in BEAST.

The majority of the positively selected sites were associated with established immune- reactive epitopes. In the Chinese poultry lineage, those were mainly located in the receptor binding subdomain (Supplementary Fig. S11). For example, positive selection was observed at sites 156, 157, and 242, which are recognized as CD8+ T- cell epitopes ^41^, and at site 87, which is associated with antibody epitope 65C6 ^42^.

Within the late-wild bird lineage, positively selected site 252 is associated with the H5_246-260_ epitope, which induced activation of T cells in chickens immunized against the HA antigen of H5 AIV ^43^. The positive selection analysis also revealed mutations (mostly in the receptor binding subdomain) that are associated with changes within the Chinese poultry lineage (D142E, H154Q /L/N, R156V/T/M/K/N/A, S157P/A, S171D/N, Q185R/K/S/G, L285V/M/I) and with the transition of the virus from the Chinese poultry lineage to the late-wild bird lineage (E142, Q154, A/T156, P157, D171, R185, V285). The amino acid states of these sites in the early-wild bird lineage (D142, N154, R156, S157, N171, Q185, L285) are mostly different to those in the late-wild bird lineage, indicating that these changes are not reversion mutations.

## Discussion

Since 2020, H5 subtype avian influenza viruses have caused outbreaks in European and Asian countries, posing a significant real threat to the poultry industry and a potential threat to public health. We reconstructed the inter-species transmission history of H5 subtype avian influenza viruses among wild birds and poultry populations with different vaccination levels. Our analysis reveals a shift from a lineage circulating within Chinese poultry to one circulating among wild birds. The wild bird lineage has frequently transmitted to unvaccinated European poultry, while the spillback of this virus from wild birds to vaccinated poultry appears to be impeded. Further, the virus lineage circulating in highly-vaccinated Chinese poultry exhibits evidence of more non-synonymous and adaptive molecular evolution in the HA gene after the date of introduction of mass poultry vaccination. The Chinese poultry lineage may have experienced more vaccine-driven selection than the other lineages. Further research is needed to determine if this selection has had any impact on the HA gene mutations present in the late-wild bird lineage.

Due to high virus prevalence, Chinese poultry, as well as Southeast Asian poultry, have been regarded as the primary reservoirs of H5 HPAIV ^44^. H5 AIV spread to other regions through bird migration ^45^ and the poultry trade ^46^, and our results reveal a 2-3- year delay between the peak of lineage dissemination from Chinese poultry and the peak of lineage introduction into European poultry. Although it has been suggested that the intercontinental spread of H5 AIV may occur within a single avian migratory cycle ^8,45, 47–50^, the delay we observe may be attributed in part to pre-existing immunity in wild birds, which may provide partial protection against infection and disease ^51–53^, leading to reduced circulation and potentially dampening large-scale outbreaks in wild birds. However, the duration of protection conferred by previous AIV infection and its impact on the epidemiology of AIV have yet to be elucidated fully ^54^. Furthermore, the relatively short generation length of many wild birds (∼ 3 years) may result in higher turnover ^55^ of serologically-naïve wild birds in nature. As the reservoir of H5 AIV shifted from Chinese poultry to wild birds, frequent migration and large-scale spatial distribution of wild birds likely facilitated inter-species transmission between wild birds and poultry populations ^45^. It is crucial that countries and regions enhance regular surveillance of avian influenza viruses in wild birds and actively and closely monitor the dynamics of virus transmission.

Following the mass poultry vaccination strategy implemented in China since 2005, the spread of H5 AIV there has been relatively well-controlled. Inter-species transmission of these viruses from or to Chinese poultry seems to be limited.

However, the Chinese poultry lineage may have experienced more antigenic evolution compared to other lineages. Mutations at amino acid positions 136, 142, 157, 172, 201, and 205 in H5 AIV HA gene have been shown to reduce reactivity to specific antibodies ^56^. Amino acids at positions 142, 172, and 205 also appear to function as immunodominant epitopes in H5 viruses ^56^. We detected positive selection at positions 142, 157, 172, and 205 in the Chinese poultry lineage but not in the wild bird lineages or the other poultry lineages. The H5N6 virus currently circulating in Chinese poultry exhibits antigenic divergence from the strains included in the commercial vaccine in China ^57, 58^, potentially leading to reduced vaccine effectiveness. Our study indicates that when vaccination is used, regular monitoring and refinement of vaccines to target emerging escape variants is necessary to respond to the emergence of novel viruses.

Whilst contemporary H5 HPAIVs are considered unlikely to acquire the ability to infect and stably circulate among the human population ^59^, there is still an urgent need to control the spread of the virus among wild birds, not only for the preservation of wildlife but also for ensuring the safety of poultry ^1^.

Our study has several limitations. First, although we obtained robust results supported by different datasets, the heterogeneous sampling rates of infected wild birds and poultry may bias ancestral reconstructions. Secondly, although there is a high vaccination rate among Chinese poultry ^27^, extracting viral lineages circulating exclusively in vaccinated poultry was not possible due to the unknown vaccination status of all sampled hosts. Thirdly, the accurate identification of ancestral host states in AIV lineages is challenging for countries with limited AIV surveillance in wild bird populations. Finally, the lack of viral genetic and surveillance data from many vaccinating/non-vaccinating countries may preclude comparative analysis of evolutionary dynamics among poultry lineages with different vaccination levels.

In conclusion, we find that vaccination in Asian poultry likely reduced the inter- species transmission of these viruses. H5 AIV in Chinese poultry, which are highly vaccinated, show evidence of greater HA gene molecular evolution and adaptation after the introduction of vaccination. Such circumstances may have increased the probability that birds susceptible to AIV belong to wild species at the interface between wild birds and poultry, leading to shifts in selection pressure on the virus. As avian influenza continues to pose significant challenges to wild and domestic animal health, our research can help inform the development of preventive measures against AIV, such as global vaccination policies.

## Method Sequence data

We collected publicly-available hemagglutinin (HA) gene sequences of H5 AIV sampled in Asia and Europe from January 1996 to February 2023 from Global Initiative on Sharing All Influenza Data (GISAID). Only sequences with available information on date and sampling location were retained for further analysis. We aligned the sequences using MAFFT v7.487 ^60^ and recombinant sequences were detected using RDP4 ^61^. We identified and removed sequences with unexpectedly high or low levels of genetic divergence given their sampling time from our datasets by estimating a maximum likelihood tree using FastTree v2.1.11 ^62^ under the automatically determined best-fit substitution model and performing a root-to-tip regression analysis in TempEst v1.5.3 ^63^.

Based on the sustained sampling efforts and the appropriate viral sample size (> 500 sequences; Supplementary Fig. S1), six Asian countries (China, Bangladesh, Japan, Korea, Indonesia, and Vietnam) were selected. All European countries were collectively treated as a single group. To focus on the viral lineages circulating in wild birds and poultry, the sequences were categorized into eight groups based on both geographic information and host type. These groups include wild birds, European poultry (without vaccination against H5 AIV), Japanese poultry (without vaccination against H5 AIV), Korean poultry (without vaccination against H5 AIV), Bangladeshi poultry (with vaccination against H5 AIV; low vaccination coverage), Indonesian poultry (with vaccination against H5 AIV; low vaccination coverage), Vietnamese poultry (with vaccination against H5 AIV; low vaccination coverage), and Chinese poultry (with vaccination against H5 AIV; high vaccination coverage).

We then downsampled these datasets in a stratified manner to create a more equitable distribution of AIV sequences between wild birds and poultry: (1) wild bird dataset: randomly selected at most 2 sequences per month per country outside China and per month per province in China, comprising 1087 gene sequences from January 1999 to January 2023; (2) European poultry dataset: randomly selecting at most 1 sequence per month per country, including 338 gene sequences from January 1997 to January 2023; (3) Japanese poultry dataset: randomly selected at most 2 sequences per month, including 72 HA gene sequences from January 2000 to January 2023; (4) Korean poultry dataset: randomly selected at most 2 sequences per month, including 72 gene sequences from October 2008 to October 2022; (5) Bangladeshi poultry dataset: randomly selected at most 1 sequence per month, including 106 HA gene sequences from January 2007 to August 2022; (6) Indonesian poultry dataset: randomly selected at most 1 sequence per month, including 121 HA gene sequences from January 2003 to March 2022; (7) Vietnamese poultry dataset: randomly selected at most 1 sequence per month, including 151 HA gene sequences from 2003 to December 2021; (8) Chinese poultry dataset: randomly selecting at most 1 sequence per month per province, including 462 gene sequences from January 1996 to March 2022.

Considering that also substantial mutations have accumulated in the PB2 gene ^64, 65^, we used a similar method to collate PB2 gene sequences of H5 AIV sampled in Asian and Europe for sensitivity analysis (see Supplementary Materials).

## Phylogenetic inference

Evolutionary histories were estimated with the Bayesian phylogenetic package BEAST v1.10.4 ^66^, using the BEAGLE ^67^ library to improve computational speed. Specifically, we employed a SRD06 substitution model ^45^, an uncorrelated lognormal relaxed clock ^45^ and Gaussian Markov random field (GMRF) Bayesian Skygrid coalescent model ^68^. We subsequently used an eight-state discrete trait analysis (DTA) implemented in BEAST 1.10.4 to infer ancestral node hosts on empirical distributions of 500 time-calibrated trees sampled from the posterior tree distributions ^69^. An asymmetric model was used for the host discrete trait, which allows different rates of lineage movement between each pair of host states ^70^. Three independent Markov chain Monte Carlo (MCMC) runs were performed for 400 million steps and logged every 20,000 steps. The first 10% of each chain was discarded as burn-in. We confirmed the convergence of all chains in Tracer v1.7.1 ^71^, ensuring the ESS was >200 for all parameters. A maximum clade credibility tree was estimated using TreeAnnotator v1.10.4 and subsequently visualized using FigTree v1.4.4 (http://tree.bio.ed.ac.uk/software/figtree) along with the R package ggtree v2.4.1 ^72^. We used the BaTS 2.0 software ^73^ to investigate the uncertainty arising from phylogenetic error (grouped by sampling location and host), which was compared to a null hypothesis that there was no association between the phylogenetic structure and traits by performing tip randomization with 1000 replicates (Supplementary Table S9).

## Evolutionary analysis within host-specific populations

To infer virus dissemination within host-specific populations, we removed AIV phylogenetic clades that were determined to represent transmissions between different species ^46^. We extracted sequences for each host-specific lineage based on the aforementioned discrete trait analysis and independently estimated the temporal phylogenies using the same substitution, clock and tree models as described above.

Strong phylogenetic temporal structure was observed in all host-specific lineages (Supplementary Fig. S12).

## Divergence analysis

To estimate site-specific synonymous and non-synonymous substitutions in the different host-specific lineages, we applied a renaissance counting ^40^ procedure implementing a codon-position specific HKY nucleotide substitution model along with an uncorrelated lognormal relaxed clock ^45^ and the Gaussian Markov random field (GMRF) Bayesian Skygrid coalescent model ^68^. Sites under positive selection were also identified using complementary approaches. Specifically, the Fast Unconstrained Bayesian AppRoximation (FUBAR) ^74^, Single Likelihood Ancestor Counting (SLAC) methods ^75^ and Fixed Effects Likelihood (FEL) methods ^75^ as implemented in Hyphy v2.5 were used ^76^.

The synonymous and nonsynonymous divergence for each host-specific lineage were calculated as the average Hamming distance between each sequenced isolate in that lineage and a reference sequence (NCBI Reference Sequence: AF144305). The estimated divergence was calculated by dividing the total number of observed differences between isolate and reference nucleotide sequence that resulted in a substitution(nonsynonymous or synonymous) by the number of possible nucleotide mutations that could result in a substitution, weighted by kappa ^77^, the transition/transversion rate ratio, which was inferred from host-specific analysis using BEAST.

## Time series analysis

The extended convergent cross mapping was applied to detect the time lags between time series variables ^78^. This analysis was performed using the R package rEDM v1.14.0 ^78^.

## Estimating rates of adaptive substitution

We employed an established population genetic method related to the McDonald- Kreitman test ^39, 79^ to estimate the number of adaptive substitutions per codon per year in HA H5 gene from the Chinese poultry lineage. We used a consensus of HA sequences from the earliest time point (sampled in 1996) as an outgroup to estimate ancestral and derived site frequencies at subsequent time points. A bootstrap analysis with 1,000 replicates was conducted to assess statistical uncertainty.

## ACKNOWLEDGMENTS

We gratefully acknowledge the authors of the originating and submitting laboratories for their crucial contributions to the generation and sharing of genome sequences and associated metadata through GISAID. This study was supported by the National Key Research and Development Program of China (2022YFC2303803); National Natural Science Foundation of China (82073616, 82204160); Beijing Advanced Innovation Program for Land Surface Science (110631111); Fundamental Research Funds for the Central Universities (2233300001); BNU-FGS Global Environmental Change Program (No.2023-GC- ZYTS-11); Research on Key Technologies of Plague Prevention and Control in Inner Mongolia Autonomous Region (2021ZD0006). J.R., and O.G.P. were supported by the UKRI GCRF One Health Poultry Hub (grant no. B/S011269/1). S.C.H. is supported by a Sir Henry Wellcome Postdoctoral Fellowship from the Wellcome Trust (220414/Z/20/Z) (https://wellcome.org/). PL acknowledges support from the European Union’s Horizon 2020 research and innovation programme (grant agreement no. 725422-ReservoirDOCS) from the European Union’s Horizon 2020 project MOOD (grant agreement no. 874850) and from the Wellcome Trust through project 206298/Z/17/Z. For the purpose of open access, the author has applied a CC BY public copyright licence to any Author Accepted Manuscript version arising from this submission. The funders had no role in study design, data collection and analysis, the decision to publish, or in preparation of the manuscript.

## DATA AND CODE AVAILABILITY

All original code and data have been deposited at Mendeley Data (https://data.mendeley.com/drafts/mmjmhc394f). The gene segment sequences are available in the GenBank (www.ncbi.nlm.nih.gov/genome/viruses/variation/flu/) and GISAID (platform.gisaid.org/) databases (https://data.mendeley.com/drafts/mmjmhc394f). Any additional information required to reanalyze the data reported in this paper is available from the lead contact upon request.

## Authors’ contributions

H.T. designed the study. H.T., O.G.P., and B.L. designed the analysis. B.L., J.R., and P.L. conducted the analyses. B.L. and Y.L. contributed and collected data. B.L. created figures. B.L., H.T., and O.G.P. wrote the initial draft. O.G.P., J.R., S.C.H., S.F., N.L., Z.W., and L.D. interpreted the data and edited the manuscript. All authors read and approved the manuscript.

