## Supplementary Material for "Association of poultry vaccination with the interspecies transmission and molecular evolution of H5 subtype avian influenza virus"

The PDF file includes:

- Supplementary Information Text
- Figs. S1 to S12
- Tables S1 to S8

**Supplementary Information Text**

**Sampling strategies for sensitivity analysis of interspecies transmission between wild birds, vaccinated poultry and unvaccinated poultry**

We performed multiple random downsamples on the original datasets in a stratified manner to obtain more robust historical dynamics of interspecies transmission among populations.

Dataset R1: 1,326 H5 AIV haemagglutinin (HA) gene sequences of domestic poultry and 793 sequences of wild birds. We performed random downsampling of the datasets, ensuring equal sampling frequency for wild birds and poultry sequences: 1) wild birds dataset (randomly selected at most 1 sequence per month per country in Europe and per month per province in China, and randomly selected at most 2 sequences per month per country among Japan, Korean, Bangladesh, Indonesia and Vietnam), comprising 793 HA gene sequences from January 1999 to January 2023; 2) European poultry dataset (randomly selected at most 1 sequence per month per country), including 338 HA gene sequences from January 1997 to January 2023; 3) Japanese poultry dataset (randomly selected at most 2 sequences per month), including 76 HA gene sequences from January 2000 to January 2023; 4) Korean poultry dataset (randomly selected at most 2 sequences per month), including 74 gene sequences from October 2008 to October 2022; 5) Bangladeshi poultry dataset (randomly selected at most 1 sequence per month), including 106 HA gene sequences from May 2007 to August 2022. 6) Indonesian poultry dataset (randomly selected at most 1 sequence per month), including 121 HA gene sequences from January 2003 to March 2022. 7) Vietnamese poultry dataset (randomly selected at most 1 sequence per month), including 151 HA gene sequences from 2003 to December 2021. 8) Chinese poultry dataset (randomly selected at most 1 sequence per month per province), including 460 HA gene sequences from January 1996 to March 2022.

Dataset R2: 1266 H5 AIV haemagglutinin (HA) gene sequences of domestic poultry and 795 sequences of wild birds. We performed random downsampling of the datasets, ensuring equal sampling frequency for wild birds and poultry sequences: 1) wild birds dataset (randomly selected at most 1 sequence per month per country in Europe and per month per province in China, and randomly selected at most 2 sequences per month per country among Japan, Korean, Bangladesh, Indonesia and Vietnam), comprising 795 HA gene sequences from January 1999 to January 2023; 2) European poultry dataset (randomly selected at most 1 sequence per month per country), including 338 HA gene sequences from January 1997 to January 2023; 3) Japanese poultry dataset (randomly selected at most 1 sequence per month), including 47 HA gene sequences from January 2000 to January 2023; 4) Korean poultry dataset (randomly selected at most 1 sequence per month), including 42 gene sequences from October 2008 to October 2022; 5) Bangladeshi poultry dataset (randomly selected at most 1 sequence per month), including 106 HA gene sequences from May 2007 to August 2022. 6) Indonesian poultry dataset (randomly selected at most 1 sequence per month), including 121 HA gene sequences from January 2003 to March 2022. 7)

Vietnamese poultry dataset (randomly selected at most 1 sequence per month), including 151 HA gene sequences from 2003 to December 2021. 8) Chinese poultry dataset (randomly selected at most 1 sequence per month), including 460 HA gene sequences from January 1996 to March 2022.

**Sampling strategies for sensitivity analysis of time lags in interspecies transmission between Chinese poultry, wild birds and European poultry**

The time lag in virus transmission was observed from Chinese poultry to wild birds, and then from wild birds to European poultry. To verify the stability of this transmission pattern, we performed sensitivity analyzes using diverse datasets. Given the singular transmission chain involving Chinese poultry, wild birds, and European poultry, the sensitivity analyses were exclusively based on virus sequences from these specific sources.

*Dataset S1*: 800 H5 AIV haemagglutinin (HA) gene sequences of domestic poultry, 850 of wild birds and 148 of environment samples. As human and other mammals are considered as almost always terminal hosts of AIVs, we only combined the sequences sampled from the environment with the main dataset (see Method section in the main text) to obtain the relatively complete phylogenies and interspecies transmission of the virus: 1) wild birds dataset (randomly selected at most 2 sequences per month per country in Europe and per month per province in China), including 850 HA gene sequences from January 1999 to January 2023; 2) European poultry dataset (randomly selected at most 1 sequence per month per country), including 338 HA gene sequences from January 1997 to January 2023; 3) Chinese poultry dataset (randomly selected at most 1 sequence per month per province), including 462 HA gene sequences from January 1996 to March 2022; 4) Environment dataset (randomly selected at most 1 sequence per month per country in Europe and province in China), including 148 HA gene sequences from January 2005 to March 2023.

*Dataset S2*: 800 H5 AIV haemagglutinin (HA) gene sequences of domestic poultry and 531 sequences of wild birds. We performed random downsampling of the datasets, ensuring equal sampling frequency for wild birds, Chinese poultry, and European poultry sequences: 1) wild birds dataset (randomly selected at most 1 sequence per month per country in Europe and per month per province in China), including 531 HA gene sequences from January 1999 to January 2023; 2) European poultry dataset (randomly selected at most 1 sequence per month per country), including 338 HA gene sequences from January 1997 to January 2023; 3) Chinese poultry dataset (randomly selected at most 1 sequence per month per province), including 462 HA gene sequences from January 1996 to March 2022.

*Dataset S3*: 800 H5 AIV haemagglutinin (HA) gene sequences of domestic poultry and 850 of wild birds. To mitigate potential sampling biases, we randomly subsampled the datasets using the same sampling strategy that was applied to the main dataset (see Method section in main text), and obtained a dataset with roughly equal numbers of sequences from wild bird and poultry: 1) wild birds dataset (randomly selected at most 2 sequences per month per country in Europe and per month per province in China), including 850 HA gene sequences from January 1999 to January 2023; 2) European poultry dataset (randomly selected at most 1 sequence per month per country), including 338 HA gene sequences from January 1997 to January 2023; 3) Chinese poultry dataset (randomly selected at most 1 sequence per month per province), including 462 HA gene sequences from January 1996 to March 2022.

**Sampling strategies for sensitivity analysis of evolutionary rate in vaccinated poultry**

Chinese poultry has relatively abundant and complete virus sequence samples, so we calculated the evolutionary rate of H5 AIV PB2 gene in wild birds and Chinese poultry.

*Dataset PB2*: 729 H5 AIV PB2 gene sequences of domestic poultry and 817 of wild birds. To mitigate potential sampling biases, we randomly subsampled the datasets using the same sampling strategy that was applied to the main dataset (see Method section in main text), and obtained a dataset with roughly equal numbers of sequences from wild birds and poultry: 1) wild birds dataset (randomly selected at most 2 sequences per month per country in Europe and per month per province in China), including 817 HA gene sequences from April 1996 to May 2023; 2) European poultry dataset (randomly selected at most 1 sequence per month per country), including 327 HA gene sequences from January 1997 to May 2023; 3) Chinese poultry dataset (randomly selected at most 1 sequence per month per province), including 402 HA gene sequences from January 1997 to March 2022.

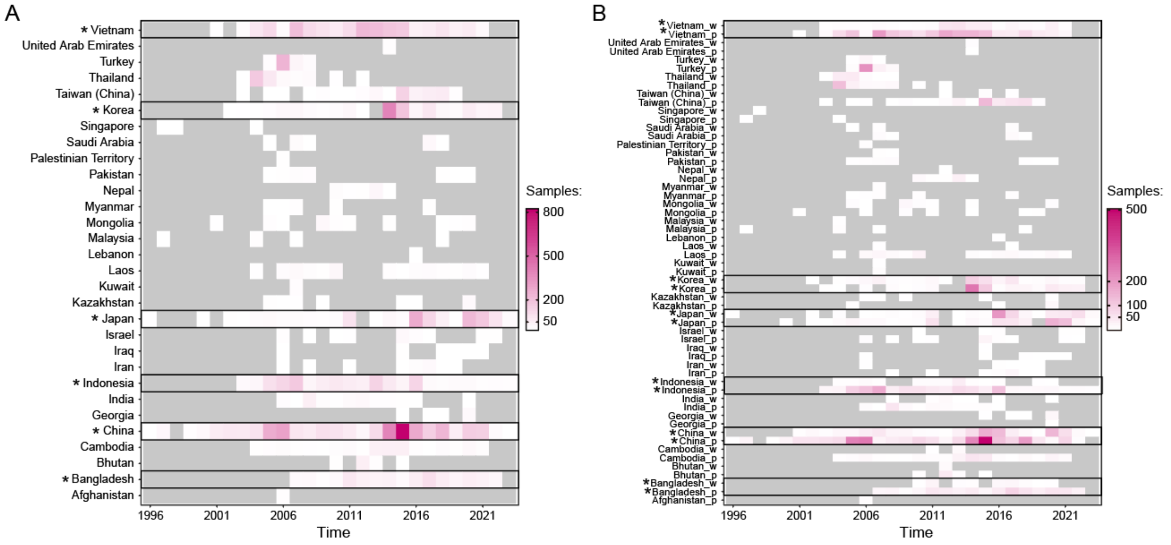


**Fig S1. Number of H5 AIV HA gene sequences sampled in Asian countries and regions.** (A) Total sample size. (B) Number of gene sequences sampled from different hosts (w: sampled from wild birds, p: sampled from poultry). Countries with continuous sampling and an adequate sample size (>500 sequences), from which data were retained for subsequent analysis, are marked with a black border and an asterisk.

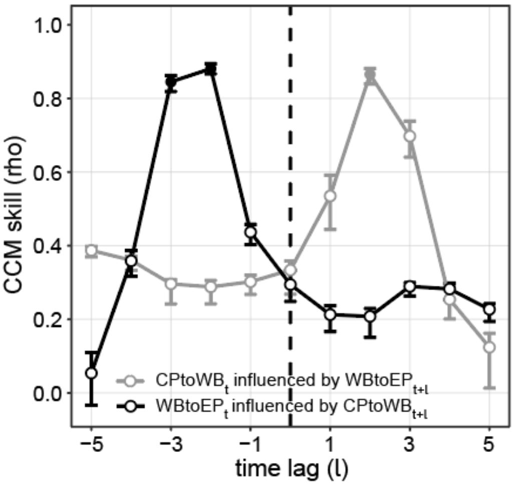


**Fig S2. Extended convergent cross mapping (CCM) analysis of interspecies transmission.** CPtoWB$_t$: the mean Markov jumps from Chinese poultry to wild bird in year t; WBtoEP$_t$: the mean Markov jumps from wild bird to European poultry in year t. We randomly selected 20 states from the converged MCMC chains and combined them into an extended chain for CCM analysis. Black line: using CPtoWB$_{t+l}$ to predict WBtoEP$_t$. The negative optimal CCM skill indicated that CPtoWB can effectively predict WBtoEP 2-3 years ahead. Gray line: using WBtoEP$_{t+l}$ to predict CPtoWB$_t$. Conversely, the positive optimal CCM skill indicated that WBtoEP can predict CPtoWB 2-3 years in the past, but implies that forecasting future CPtoWB from WBtoEP is not feasible.

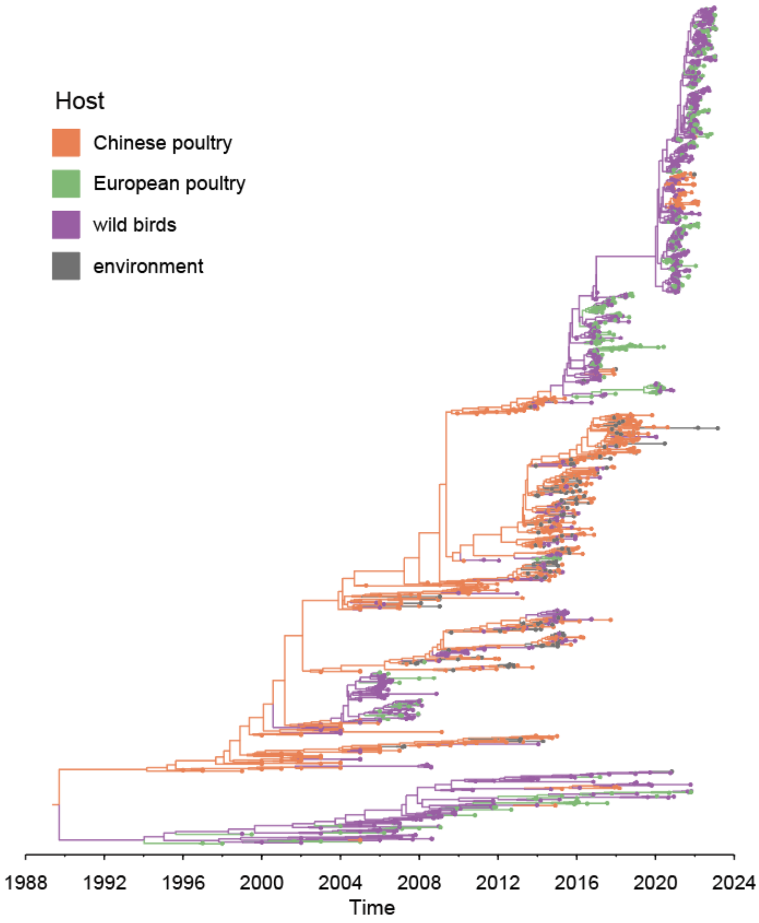


**Fig S3. The maximum clade credibility tree of H5 AIV HA gene sampled from 1996 to 2023.** The tips are colored based on their host states, while internal branches are colored according to ancestral states inferred using the asymmetric discrete phylogenetic model with Bayesian Stochastic Search Variable Selection (purple: wild birds; orange: Chinese poultry; light purple: European poultry; gray: environment). The phylogeny is inferred from *Dataset S1*.

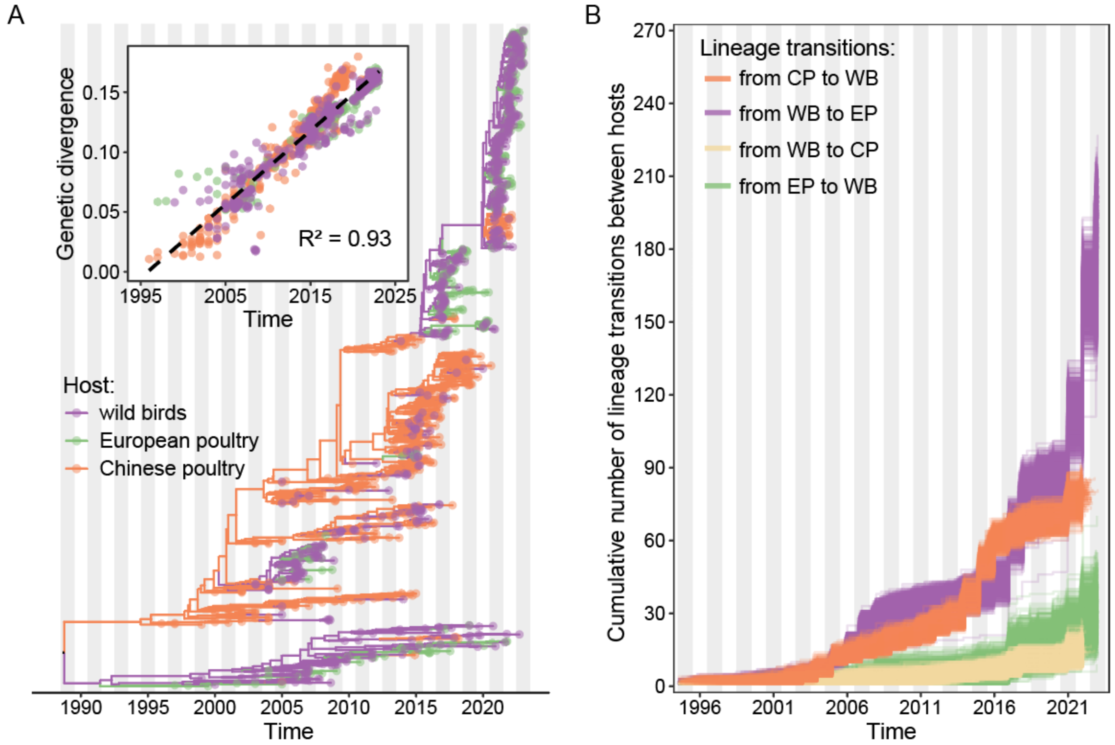


**Fig S4. Dynamic lineage transmissions of H5 AIV HA gene among three host populations inferred from Dataset S1.** (A) The maximum clade credibility tree of H5 AIV HA gene sampled from 1996 to 2022. The tips are colored based on their host states, while internal branches are colored according to ancestral states inferred using the asymmetric discrete phylogenetic model with Bayesian Stochastic Search Variable Selection (purple: wild birds; orange: Chinese poultry; light purple: European poultry). Inset: a root-to-tip regression of genetic divergence against the dates of sample collection. (B) The cumulative lineage transitions of the HA gene between the three host states were summarized from the posterior samples of an asymmetric discrete phylogenetic analysis.

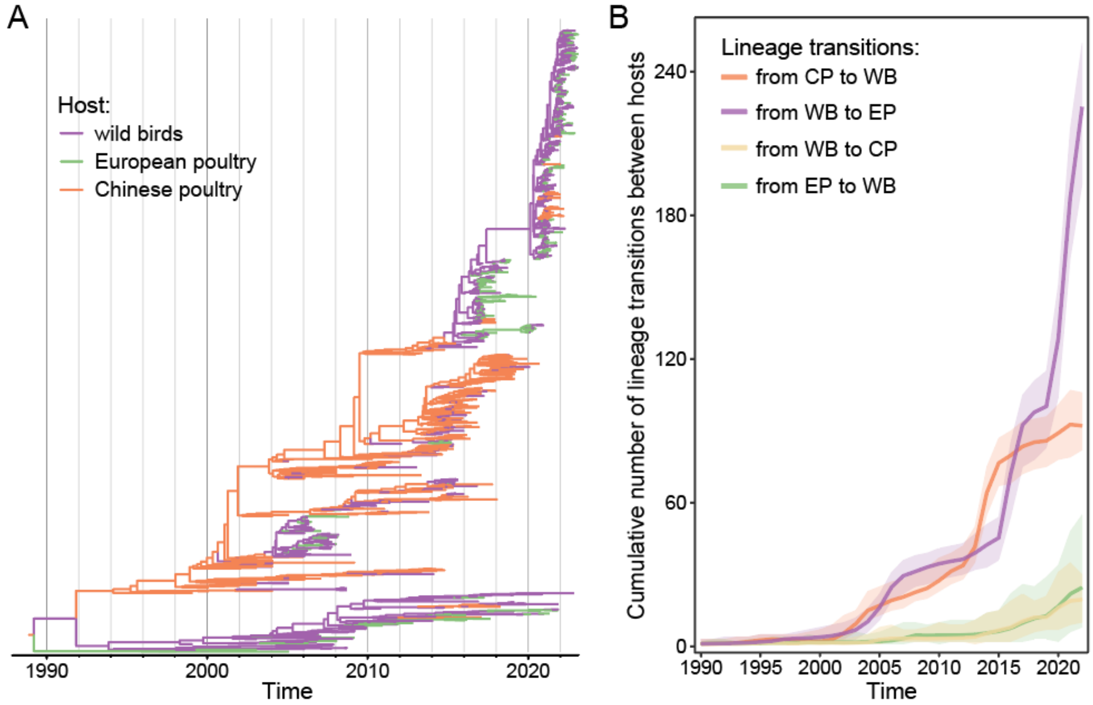


**Fig S5. Dynamic lineage transmissions of H5 AIV HA gene among three host populations inferred from Dataset S3.** (A) The maximum clade credibility tree of H5 AIV HA gene sampled from 1996 to 2022. The tips are colored based on their host states, while internal branches are colored according to ancestral states inferred using the asymmetric discrete phylogenetic model with Bayesian Stochastic Search Variable Selection (purple: wild birds; orange: Chinese poultry; light purple: European poultry). (B) The cumulative lineage transitions of the HA gene between the three host states were summarized from the posterior samples of an asymmetric discrete phylogenetic analysis. The solid coloured lines represent the annual mean values of the cumulative lineage transitions and the shaded areas show the 95% highest posterior density credible intervals of that estimate.

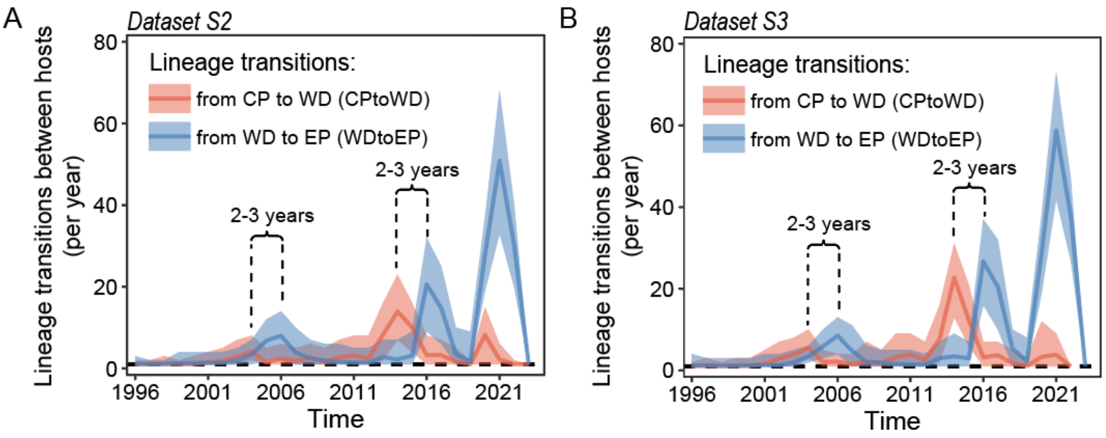


**Fig S6. Time series of the annual mean lineage transitions of H5 AIV HA gene between wild birds (WB), Chinese poultry (CP) and European poultry (EP).** Panel (A) displays results derived from Dataset S2, while Panel (B) derived from Dataset S3.

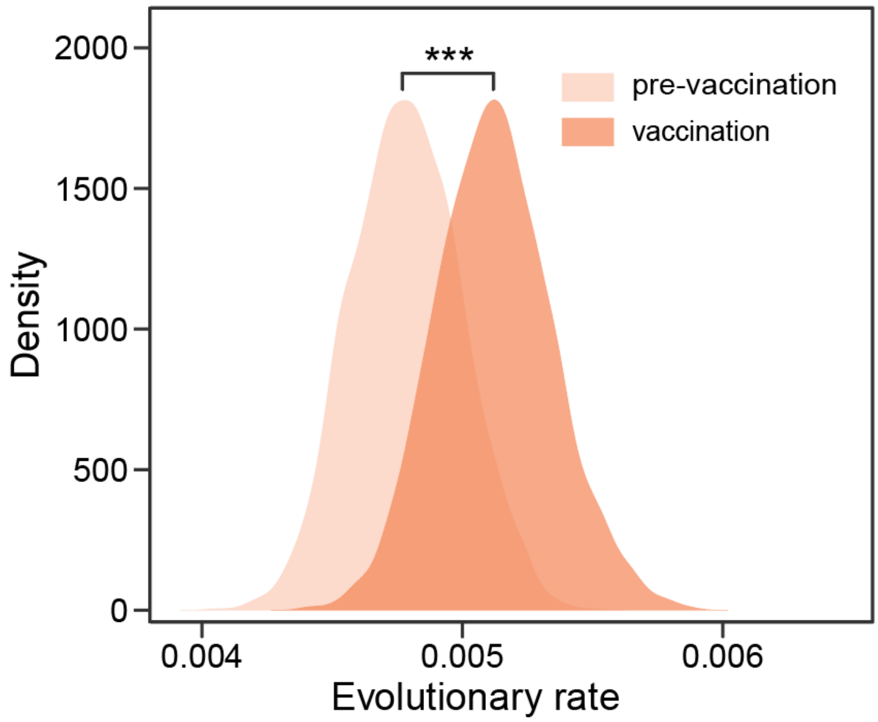


**Fig S7. Evolutionary rate (subs/site/year) of the Chinese poultry lineage in pre-vaccination era and vaccination era.** A time-dependent rate (TDR) model was implemented.

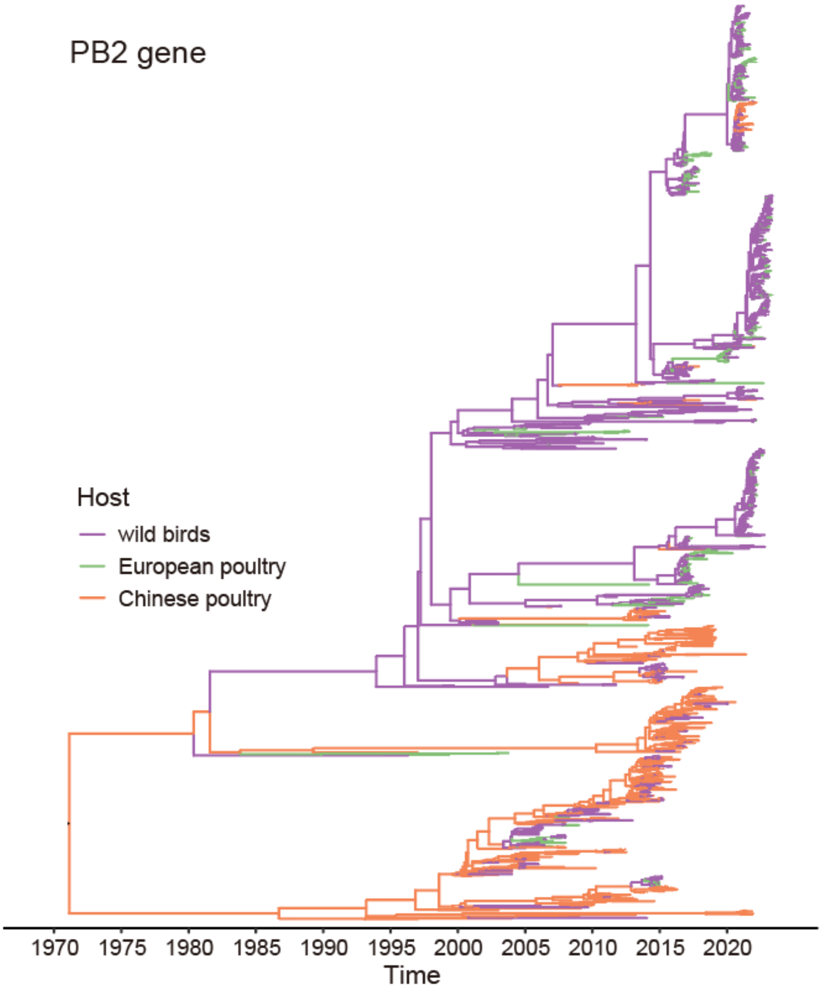


**Fig S8. Maximum clade credibility tree of the PB2 gene of H5 AIV sampled from 1996 to 2023 in China and Europe.** The tips are colored based on their host states (purple: wild birds; orange: Chinese poultry; green: European poultry), while internal branches are colored according to ancestral states inferred using the asymmetric discrete phylogenetic model with Bayesian Stochastic Search Variable Selection (purple: wild birds; orange: Chinese poultry; green: European poultry).

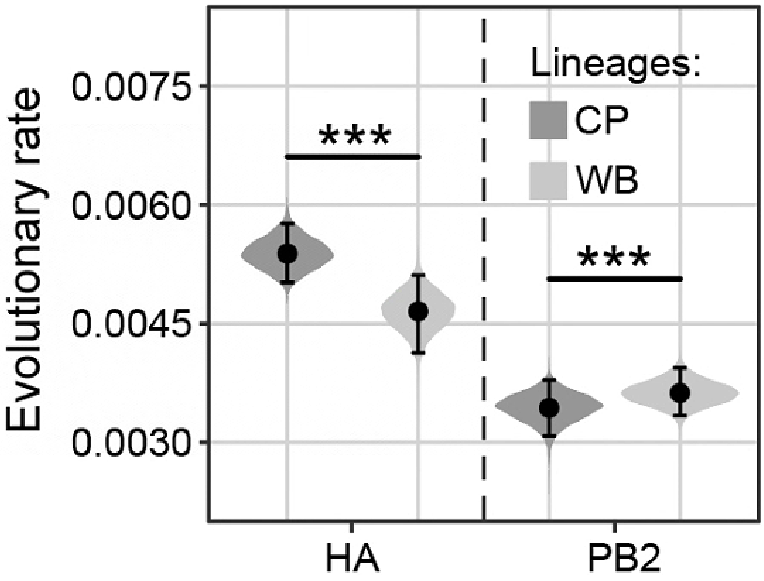


**Fig S9. The estimated substitution rates (subs/site/year) of HA gene and PB2 gene of Chinese poultry and wild bird lineages.** For the PB2 gene, there are two host-specific lineages: the Chinese poultry lineage (CP) and the wild bird lineage (WB). The lineages of HA gene were categorized into two lineages: the Chinese poultry lineage (CP), the wild bird lineage (WB, consisting of early-wild bird lineage and late-wild bird lineage). The accompanying dot and the whisker plots indicate the mean and the 95% highest posterior density credible intervals for these estimates. We used Wilcoxon tests to assess differences in pairwise substitution rates between lineages, with statistical significance levels of $p < 0.05$ (*), $p < 0.01$ (**), and $p < 0.001$ (***) denoting statistically significant, highly significant, and extremely significant differences, respectively.

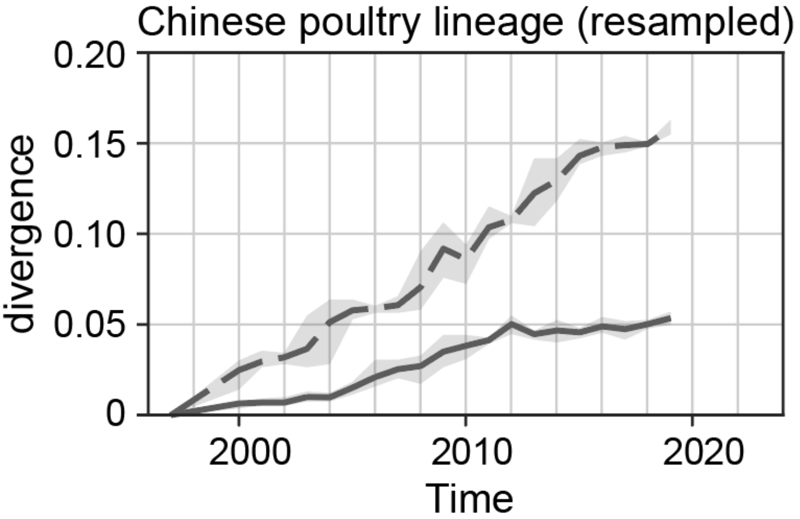


**Fig S10. Nonsynonymous (solid lines) and synonymous (dashed lines) divergence of the HA gene of the Chinese poultry lineage through time.** The Chinese poultry lineage was randomly resampled to include at most 3 sequences per year. Divergences were computed using 1-year sliding windows. Shaded region shows 95% confidence intervals.

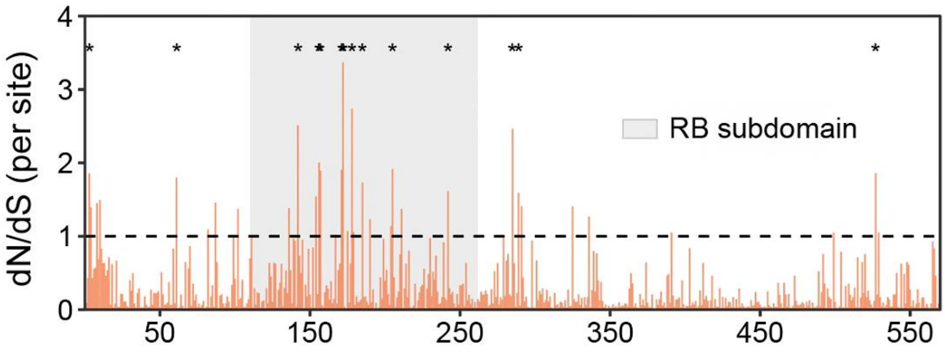


**Fig S11. Mean nonsynonymous/synonymous divergence (dN/dS) of each site in H5 AIV HA gene of the Chinese poultry lineage.** The sites identified as experiencing significant positive selection through the renaissance counting method are marked with an asterisk. The horizontal dashed line indicates the dN/dS ratio of 1, and the grey shaded regions indicate the receptor binding subdomain sites.

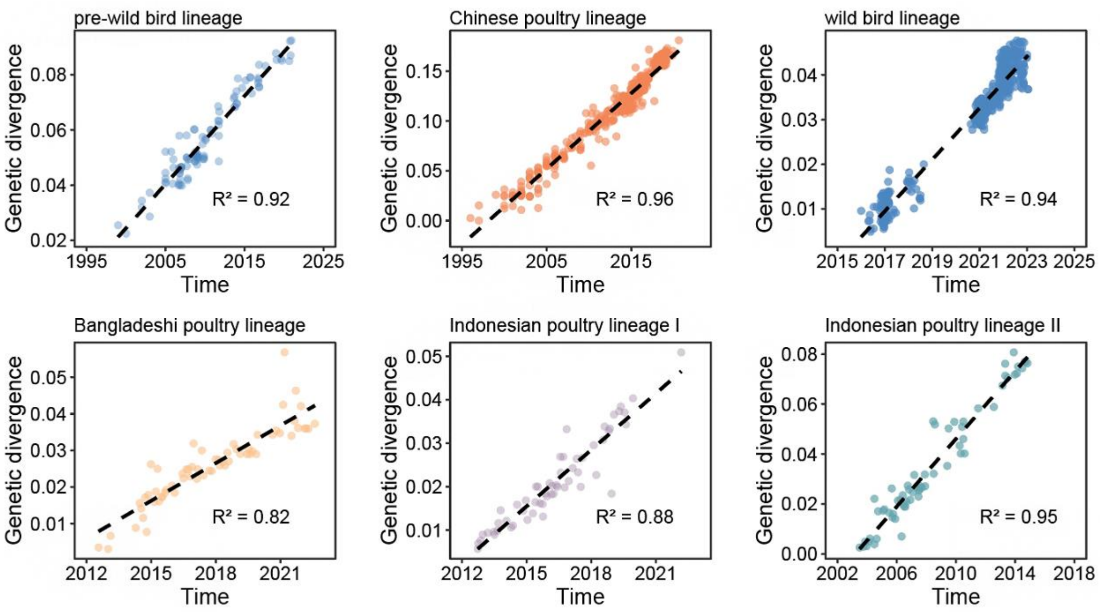


**Fig S12. Temporal signal of genetic divergence for host-specific lineages.**

**Table S1. The posterior support of each host branch in different datasets.**

| Dataset | Branch | Host. prob |
|---|---|---|
| main text | Chinese poultry section | 0.9882 |
| | Early-wild bird section | 0.7902 |
| | Late-wild bird section | 0.9822 |
| | Bangladeshi poultry lineage | 1 |
| | Indonesian poultry lineage I | 0.9882 |
| | Indonesian poultry lineage II | 1 |
| Dataset S1 | Chinese poultry section | 0.9947 |
| | Early-wild bird section | 0.824 |
| | Late-wild bird section | 0.9921 |
| Dataset S2 | Chinese poultry section | 1 |
| | Early-wild bird section | 0.8275 |
| | Late-wild bird section | 0.9935 |
| Dataset S3 | Chinese poultry section | 0.9987 |
| | Early-wild bird section | 0.8507 |
| | Late-wild bird section | 0.9934 |

**Table S2. The result of null hypothesis tests.**

| Statistic | observed | | | null | | | significance |
|---|---|---|---|---|---|---|---|
| | mean | lower 95% CI | upper 95% CI | mean | lower 95% CI | upper 95% CI | |
| AI | 65.87 | 62.96 | 69.83 | 161.00 | 155.66 | 166.03 | 0 |
| PS | 488.23 | 480 | 496 | 1051.65 | 1033.30 | 1071.21 | 0 |
| MC (state 0) | 29 | 29 | 29 | 4.81 | 4.04 | 6.12 | P<= 0.001 |
| MC (state 1) | 9.67 | 9 | 12 | 3.31 | 2.69 | 4.15 | P<= 0.001 |
| MC (state 2) | 16.85 | 16 | 22 | 6.12 | 5.06 | 8.02 | P<= 0.001 |

*HPD CIs = highest posterior density confidence intervals (credible sets).

*state 0: vaccinated poultry; state 1: unvaccinated poultry; state 2: wild birds.

**Table S3. Number of cumulative lineage transitions between wild birds and poultry populations with different vaccination statuses.**

| Dataset | Lineage transitions between hosts | Mean | 95% HPD |
|---|---|---|---|
| Dataset R1 | wild birds to vaccinated poultry populations | 31 | (22, 39) |
| | vaccinated poultry populations to wild birds | 127 | (118, 136) |
| | wild birds to unvaccinated poultry populations | 263 | (245, 280) |
| | unvaccinated poultry populations to wild birds | 53 | (38, 70) |
| Dataset R2 | wild birds to vaccinated poultry populations | 33 | (24, 41) |
| | vaccinated poultry populations to wild birds | 128 | (118, 136) |
| | wild birds to unvaccinated poultry populations | 243 | (223, 259) |
| | unvaccinated poultry populations to wild birds | 51 | (34, 66) |

**Table S4. Segmented linear regression of nonsynonymous divergence in the Chinese poultry lineage from 1997 to 2019.**

| (nonsynonymous divergence ~ year) | Parameter | Estimate | Standard Error | 95%CI |
|---|---|---|---|---|
| | Intercept | -3.3933 | 0.5621 | |
| 1996-2004 | $slope_1$ | 0.0017 | 0.0002 | (0.0011, 0.0023) |
| 2005-2010 | $slope_2$ | 0.0046 | 0.0002 | (0.0041, 0.0052) |
| 2010-2022 | $slope_3$ | 0.0013 | 0.0002 | (0.0010, 0.0017) |

*adjusted R-squared: 0.9948; multiple R-Squared: 0.9961; residual standard error: 0.001294.

**Table S5. Selection pressure in the HA gene of different host populations.**

| Host | cNrate (95% HPD) | cSrate (95% HPD) | dnds (95% HPD) |
|---|---|---|---|
| Chinese poultry | 0.001567 (0.001564, 0.001571) | 0.002752 (0.002745, 0.002758) | 0.2416 (0.2403, 0.2429) |
| European poultry | 0.001091 (0.001081, 0.001102) | 0.002890 (0.002874, 0.002906) | 0.1588 (0.1565, 0.1610) |
| wild birds | 0.001130 (0.001125, 0.001135) | 0.002820 (0.002810, 0.002829) | 0.1705 (0.1690, 0.1720) |

*cNrate, the absolute nonsynonymous rate (per site per year); cSrate, the absolute synonymous rate (per site per year).

**Table S6. Positively-selected sites in the HA gene in resampled-Chinese poultry lineage.**

| **Lineage** | **Methods** | | | |
|---|---|---|---|---|
| | **RC** | **FEL ($p$ < 0.1)** | **SLAC ($p$ < 0.1)** | **FUBAR** (PP **> 0.9)** |
| Resampled-Chinese poultry lineage | 61, 87, 136, 142, 154, 156, 157, 172, 178, 190, 205, 285, 289 | 61, 87*, 131, 142, 154*, 156, 157*, 172*, 190, 205, 216, 225 | 87, 154*, 157, 172, 205 | 87*, 131, 154*, 156*, 157*, 172, 205* |

*Asterisks mark sites inferred to be under positively selected with posterior probability (PP) >0.95 and <0.05. FEL, Fixed Effects Likelihood. SLAC, Single Likelihood Ancestor Counting. FUBAR, Fast Unconstrained Bayesian AppRoximation. RC, renaissance counting method implemented in BEAST.

**Table S7. Positively-selected sites in the HA gene among different host lineages inferred from *Dataset S1*.**

| **Lineage** | **FEL ($p < 0.1$)** | **SLAC ($p < 0.1$)** | **FUBAR (PP > 0.9)** |
|---|---|---|---|
| Chinese poultry lineage | 3, 8*, 87*, 111*, 142, 145, 154*, 157, 171*, 172*, 185, 285, 291*, 325 | 3, 142*, 143, 154*, 157*, 172*, 185*, 285* | 142*, 154*, 157*, 171, 285* |
| late-wild bird lineage | 507 | - | - |
| early-wild bird lineage | 102*, 170*, 171 | 102, 107 | 102*, 170*, 171* |

*Asterisks mark sites inferred to be under positively selected with posterior probability (PP) >0.95 and <0.05. FEL, Fixed Effects Likelihood. SLAC, Single Likelihood Ancestor Counting. FUBAR, Fast Unconstrained Bayesian AppRoximation.

**Table S8. Positively-selected sites in the HA gene among different host lineages inferred from *Dataset S2*.**

| **Lineage** | **FEL ($p$ < 0.1)** | **SLAC ($p$ < 0.1)** | **FUBAR (PP > 0.9)** |
|---|---|---|---|
| Chinese poultry lineage | 3, 8*, 87*, 111*, 142, 145, 154*, 157, 171*, 172*, 185, 285, 291*, 325 | 3, 8, 87, 111, 142, 145*, 154*, 157, 171*, 172*, 185*, 205, 338* | 87, 154*, 157, 171*, 172 |
| late-wild bird lineage | 7, 185, 204, 509 | - | - |
| early-wild bird lineage | 102*, 113, 170*, 171 | 102*, 170 | 102*, 170*, 113, 171 |

*Asterisks mark sites inferred to be under positively selected with posterior probability (PP) >0.95 and <0.05. FEL, Fixed Effects Likelihood. SLAC, Single Likelihood Ancestor Counting. FUBAR, Fast Unconstrained Bayesian AppRoximation.

**Table S9. The association between phylogeny and sampling population.**

| Statistic | observed | | | null | | | significance |
|---|---|---|---|---|---|---|---|
| | mean | lower 95% HPD CIs | upper 95% HPD CIs | mean | lower 95% HPD CIs | upper 95% HPD CIs | |
| AI | 69.38 | 66.46 | 73.49 | 187.94 | 183.38 | 192.58 | *P*=0 |
| PS | 529.69 | 521 | 538 | 1213.40 | 1199.97 | 1226.98 | *P*=0 |
| MC (state 0) | 29 | 29 | 29 | 1.79 | 1.16 | 2.13 | *P*<=0.001 |
| MC (state 1) | 14.22 | 14 | 15 | 3.24 | 2.55 | 4.07 | *P*<=0.001 |
| MC (state 2) | 9.65 | 9 | 12 | 2.71 | 2.08 | 3.46 | *P*<=0.001 |
| MC (state 3) | 7.90 | 6 | 10 | 1.88 | 1.27 | 2.17 | *P*<=0.001 |
| MC (state 4) | 4.18 | 4 | 5 | 1.49 | 1 | 2 | *P*<=0.001 |
| MC (state 5) | 6.75 | 6 | 9 | 1.47 | 1 | 2 | *P*<=0.001 |
| MC (state 6) | 13.64 | 9 | 19 | 2.03 | 1.59 | 2.54 | *P*<=0.001 |
| MC (state 7) | 16.85 | 16 | 22 | 6.12 | 5.07 | 8 | *P*<=0.001 |

*HPD CIs = highest posterior density confidence intervals (credible sets).

*state 0: Bangladeshi poultry, state 1: Chinese poultry, state 2: European poultry, state 3: Indonesian poultry, state 4: Japanese poultry, state 5: Korean poultry, state 6: Vietnamese poultry, state 7: wild birds.
